## Supplemental Figures for "Chronic stress impairs autoinhibition in neurons of the locus coeruleus to increase asparagine endopeptidase activity"

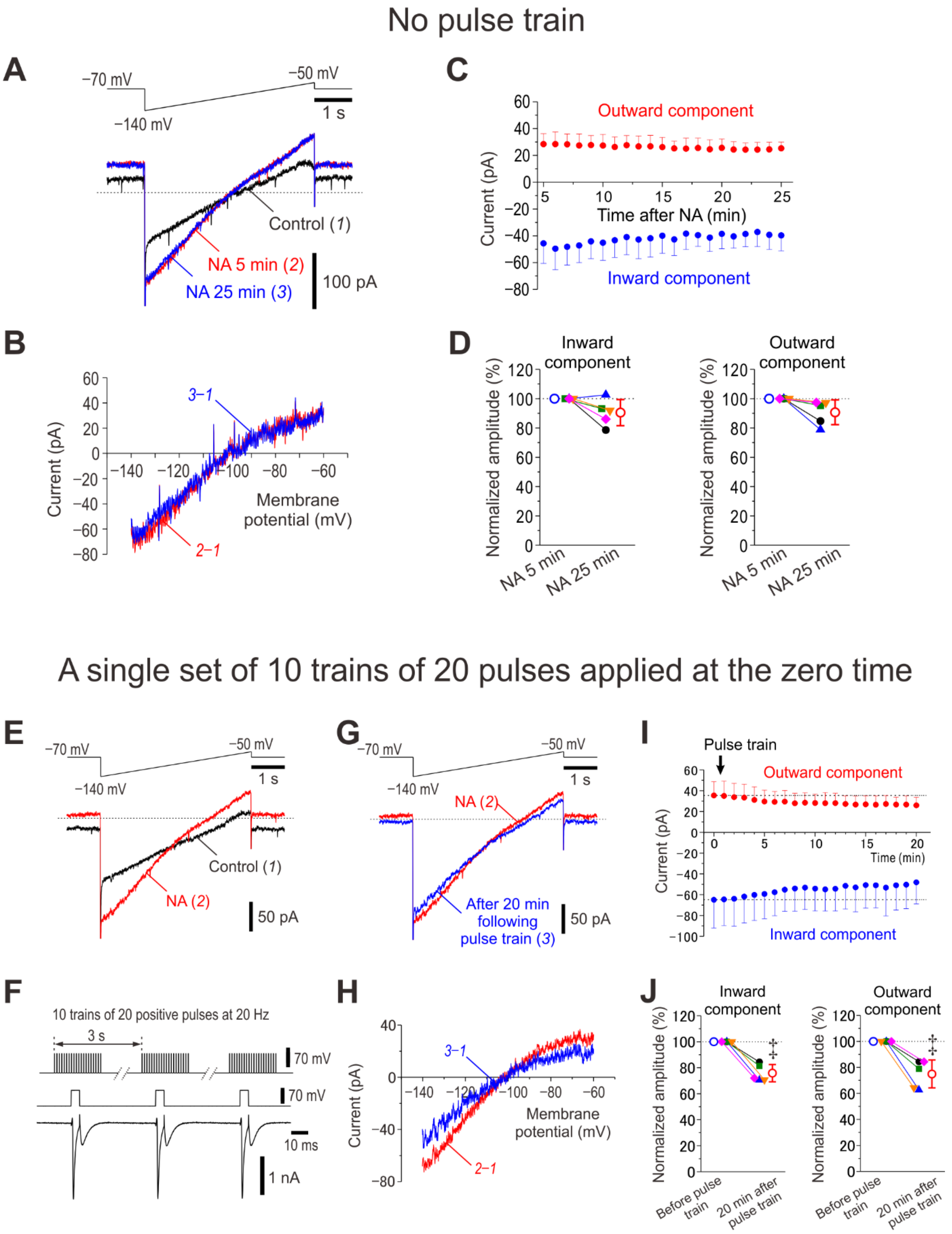


**Figure S1. NA-induced GIRK currents show no apparent agonist-dependent rundown but show a moderate rundown following application of a train of positive voltage pulses.**

(**A**) Upper panel, Ramp command pulse applied every minute. Lower panel, Superimposed current traces obtained under control condition (1), in response to 5th ramp pulse (2) and 25th ramp pulse (3) in the presence of 100 μM NA. (**B**) The I-V relationship of NA-induced GIRK currents obtained by subtraction of currents recorded under control condition from that recorded in response to the 5th ramp pulse (red trace, 2–1) and that obtained by subtraction of the control current from that recorded in response to the 25th ramp pulse (blue trace, 3–1). (**C**) Plotting of amplitudes of inward components at –130 mV (blue circles) and outward components at –60 mV (red circles) against time. (**D**) Pooled data showing no appreciable changes in normalized amplitudes of inward components at –130 mV and those of outward components at –60 mV during 20 times repetition of ramp pulses (n = 5). Inward component, paired *t*-test, $p = 0.075$; outward component, paired *t*-test, $p = 0.065$. (**E**) Upper panel, Ramp command pulse applied every minute. Lower panel, Superimposed current traces obtained under control condition (1) and in response to 5th ramp pulse in the presence of NA (2). (**F**) Upper panel, Ten trains of 20 positive pulses (5 ms duration to 0 mV at 20 Hz) at an inter-spike interval of 2 s (one pulse train) were applied only once in the presence of extracellular 30 mM TEA and intracellular 0.2 mM EGTA. Lower panel, representative $Na^+$ current followed by small $Ca^{2+}$ current in response to positive pulses. (**G**) Upper panel, Following the train of positive pulses, ramp command pulse applied every minute. Lower panel, Superimposed current traces obtained in response to 5th ramp pulse in the presence of NA (2) and 20 min following one pulse train in the presence of NA (3). (**H**) The I-V relationship of NA-induced GIRK currents obtained by subtraction of currents recorded under control condition from that recorded in response to the 5th ramp pulse in the presence of NA (red trace, 2–1) and from that recorded 20 min following one pulse train in the presence of NA (blue trace, 3–1). (**I**) Plotting of amplitudes of inward components at –130 mV (filled blue circles) and outward components at –60 mV (filled red circles) against time (n = 5). (**J**) Pooled data showing significant decreases in normalized amplitudes of inward components at –130 mV and those of outward components at –60 mV before and after 20 times repetition of ramp pulses in the presence of NA (n = 5). Inward component, paired *t*-test, $^{‡}p = 0.001$. Outward component, paired *t*-test, $^{‡}p = 0.001$.

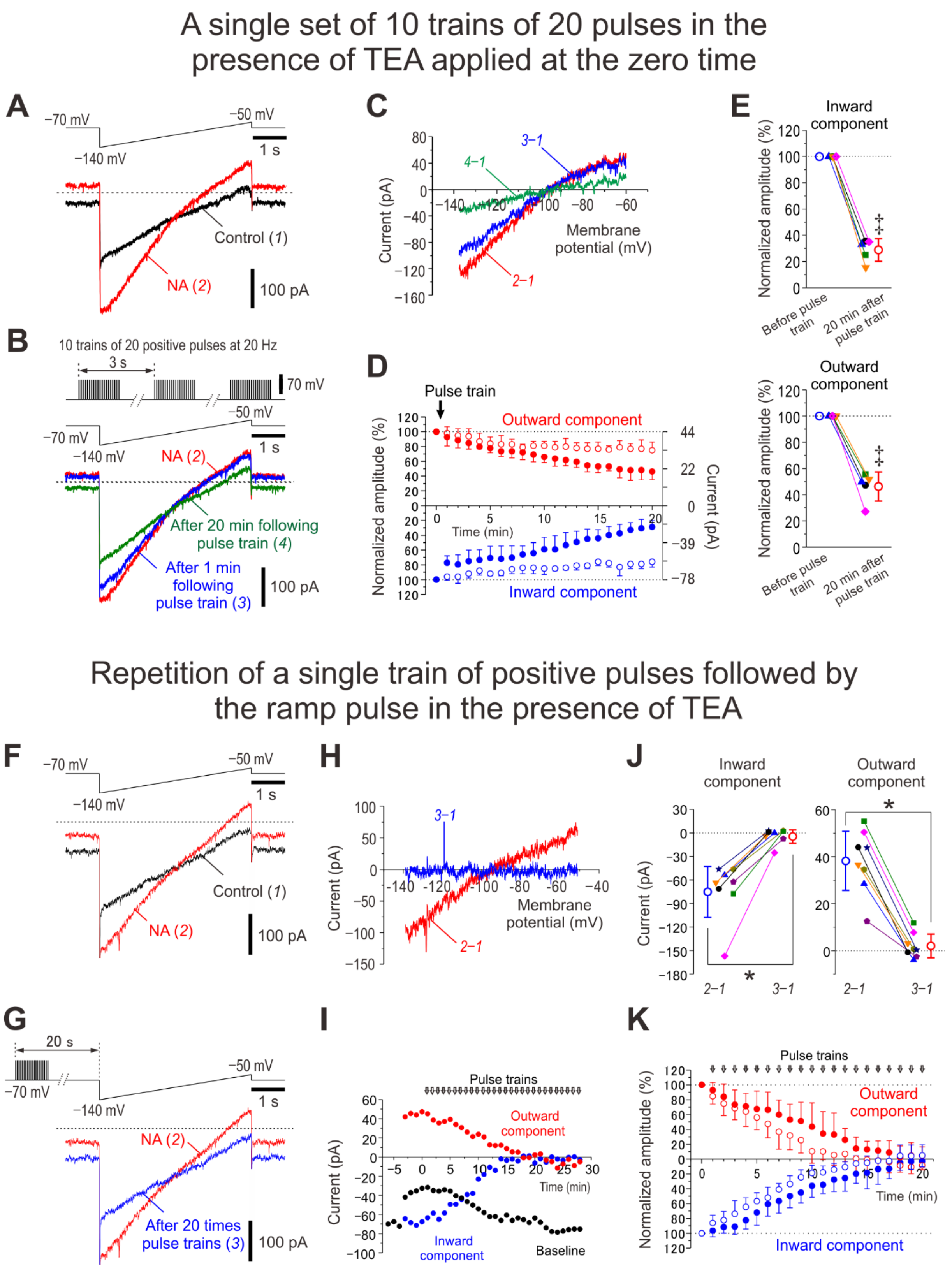


**Figure S2. Differential rundown of NA-induced GIRK currents following application of various types of positive voltage pulse trains in the presence of TEA**

(**A**) Upper panel, Ramp command pulse applied every minute. Lower panel, Superimposed current traces obtained under control condition (1) and in response to 5th ramp pulse in the presence of NA (2). (**B**) Upper panel, Ten trains of 20 positive pulses (5 ms duration to 0 mV at 20 Hz) at an inter-spike interval of 2 s (one pulse train) were applied only once in the presence of extracellular 30 mM TEA and intracellular 0.2 mM EGTA. Following one pulse train, ramp command pulses were applied every minute. Lower panel, Superimposed current traces obtained in response to 5th ramp pulse in the presence of NA (2), 1 min following pulse trains in the presence of NA and TEA (3) and 20 min following pulse trains in the presence of NA and TEA (4). (**C**) The I-V relationship of NA-induced GIRK currents obtained by subtraction of currents recorded under control condition from that recorded in response to the 5th ramp pulse in the presence of NA (red trace, 2–1), from that recorded 1 min following one pulse train in the presence of NA and TEA (blue trace, 3–1), and from that recorded 20 min following one pulse train in the presence of NA and TEA (green trace, 4–1). (**D**) Plotting of normalized amplitudes (left vertical axis) of inward components at –130 mV (filled blue circles) and outward components at –60 mV (filled red circles) against time (n = 5). Amplitudes of GIRK-I were normalized to the values obtained before applying pulse trains. Right vertical axis refers to the original amplitudes. Open blue and red circles represent the normalized amplitudes of GIRK-I, the original amplitudes of which are shown in Suppl. Fig. 1**I**. (**E**) Pooled data showing appreciable changes in normalized amplitudes of inward components at –130 mV and those of outward components at –60 mV before and after 20 times repetition of ramp pulses in the presence of NA and TEA (n = 5). Inward component, paired *t*-test, $^{‡}p < 0.001$; outward component, paired *t*-test, $^{‡}p < 0.001$. (**F**) Upper panel, Ramp command pulse. Lower panel, Superimposed current traces obtained under control condition (1) and after application of 100 μM NA for 5 min (2). (**G**) Upper panel, A combined command pulse applied every minute; one train of 20 positive pulses (5 ms duration to 0 mV at 20 Hz) in the presence of extracellular 30 mM TEA and intracellular 0.2 mM EGTA, which was followed by the ramp pulse after an interval of 19 s. Lower panel, Superimposed current traces obtained after application of NA for 5 min (2) and in response to application of the 20th combined pulse in the presence of NA and TEA (3). (**H**) The I-V relationship of NA-induced GIRK currents obtained by subtraction of currents recorded under control condition from that recorded after application of NA for 5 min (red trace, 2–1) and that obtained by subtraction of the control current from that recorded in response to application of the 20th combined pulse in the presence of NA and TEA (blue trace, 3–1). (**I**) Plotting of amplitudes of inward components at –130 mV (blue circles), outward components at –60 mV (red circles) and baseline currents (black circles) against time. (**J**) Pooled data showing decreases in amplitudes of inward components at –130 mV and those of outward components at –60 mV at respective conditions (2–1 and 3–1) (n = 8). Inward component, paired *t*-test, $^{‡}p < 0.001$; outward component, paired *t*-test, $^{‡}p < 0.001$. (**K**) Plotting of normalized amplitudes of inward components at –130 mV (filled blue circles) and outward components at –60 mV (filled red circles) against time before and during application of positive pulse trains in the presence of NA and TEA (n = 8). The amplitudes of inward components at –130 mV and those of outward components at –60 mV were normalized by those recorded in the presence of NA before applying positive pulse trains. Open blue and red circles represent results obtained in Fig. 2**E**.

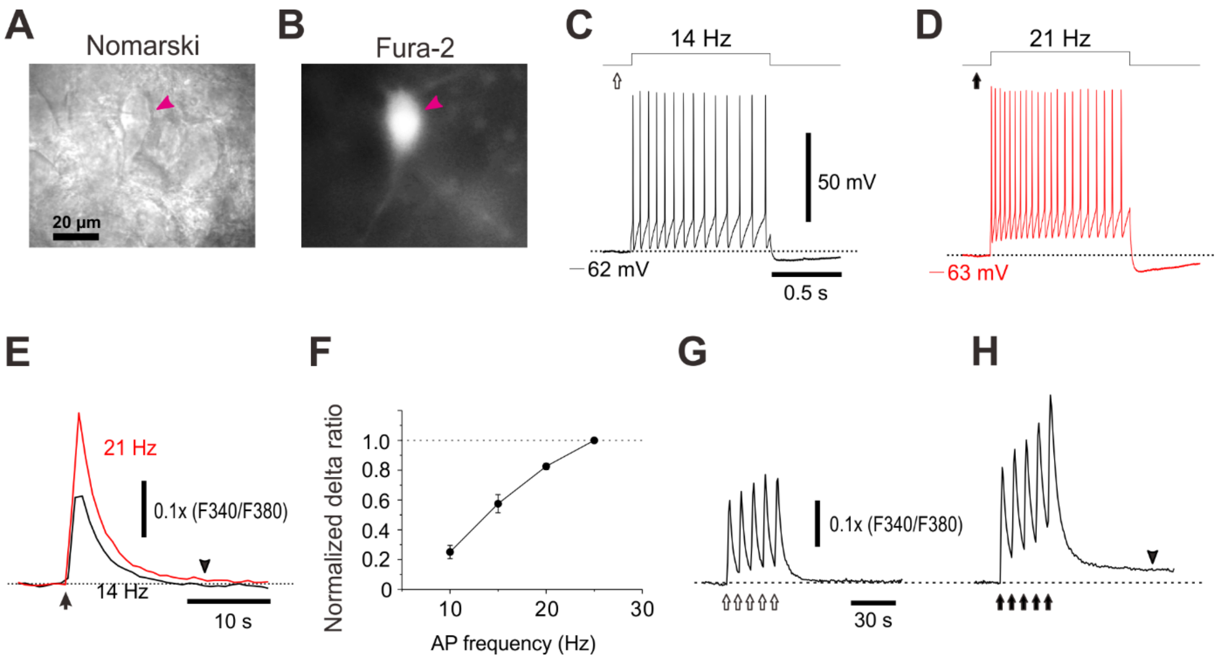


**Figure S3. $Ca^{2+}$ transients in response to current pulse injections in LC neurons.**
(**A, B**) Nomarski and fura-2 images of a fura-2 loaded LC neuron (arrowhead). (**C, D**) Spike trains at 14 and 21 Hz evoked in response to depolarizing current pulses. The maximal firing frequencies of LC neurons in response to depolarizing current pulses were usually lower than 20-25 Hz (n > 40). (**E**) $Ca^{2+}$ transients in response to current pulse injections which induce spike trains at 14 and 21 Hz in the same LC neuron. Note a long-lasting tail of $Ca^{2+}$ transient (downward arrowheads) in response to stronger activation (21 Hz) of LC neurons. (**F**) Relationship between action potential frequency and the peak amplitude of $Ca^{2+}$ transients (n = 5). (**G, H**) Differential summation of $Ca^{2+}$ transients induced by the two different sets of five trains of spike firings at 14-16 Hz and at 19-21 Hz, respectively, evoked by two different current pulses with intensities of 75 and 100 pA. Note an emergence of long-lasting slow $Ca^{2+}$ transients (downward arrowheads) in response to stronger activation (19-21 Hz) of LC neurons, suggesting an involvement of CICR in the generation of such a long lasting slow $Ca^{2+}$ transients.

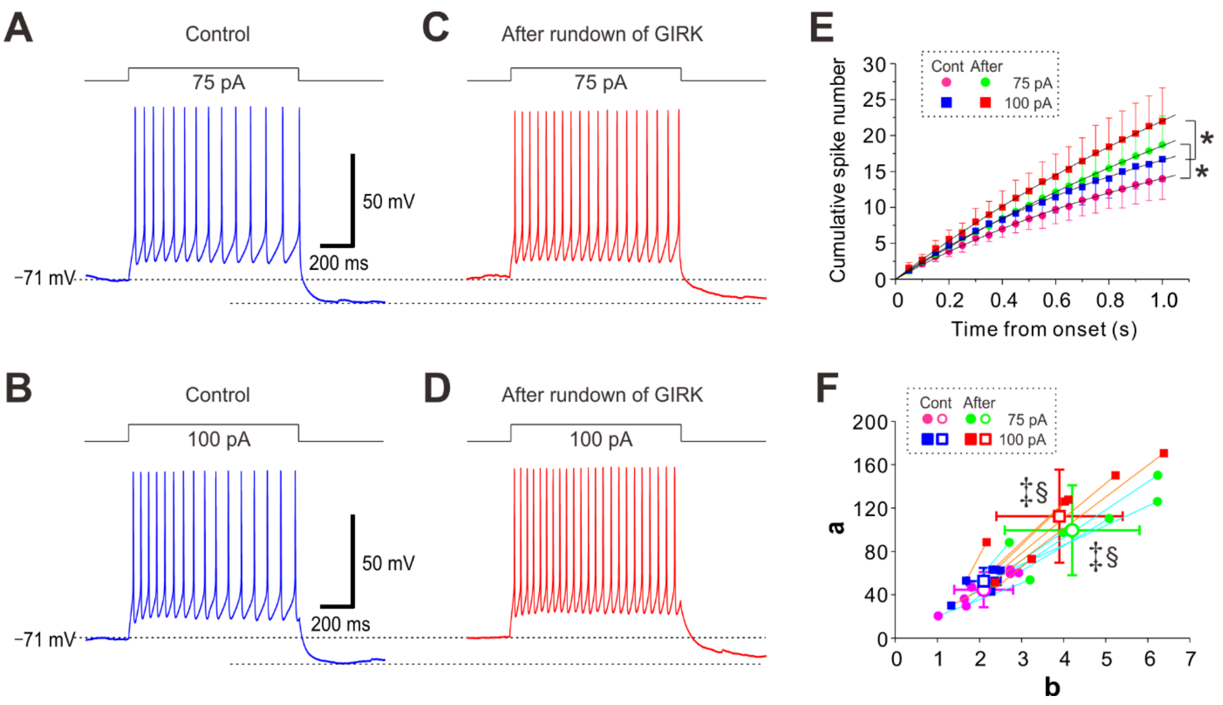


**Figure S4. Weakening of spike-frequency adaptation and decrease in pulse-AHP accompany GIRK rundown.**
(**A-D**) Sample traces of spike trains induced in an LC neuron evoked by current pulses at 75 and 100 pA under control conditions and after rundown of GIRK-I as shown in Fig. 4**A-F**. Note the abolishment of spike-frequency adaptation and reduction of pulse-AHP after rundown of GIRK-I. (**E**) Plotting of the cumulative spike numbers vs the elapsed time during the current pulses every 50 ms obtained under control condition (pink circles: 75 pA; blue squares: 100 pA) and those obtained after rundown of GIRK-I (green circles: 75 pA; red squares: 100 pA). 75 pA current pulse, two-way RM ANOVA, $^{*}p = 0.002$; 100 pA current pulse, two-way RM ANOVA, $^{*}p = 0.004$ (n = 7). (**F**) Plotting of the saturation level (***a***) vs the half saturation constant (***b***), which were measured by curve fitting to the data points in **E**. The values of ***a*** and ***b*** obtained after rundown of GIRK-I (green circles, 75 pA; red squares, 100 pA) were significantly larger than those obtained under control condition (pink circles, 75 pA; blue squares, 100 pA) (75 pA current pulse, paired t-test, ***a*** and ***b***, $^{‡}p = 0.002$ and $^{‡}p = 0.002$, respectively; 100 pA current pulse, paired t-test, ***a*** and ***b***, $^{‡}p = 0.004$ and $^{‡}p = 0.007$, respectively), and there was a significant difference in the relationship between ***a*** and ***b*** (75 pA current pulse, Wilk's lambda: $^{§}p = 0.028$; 100 pA current pulse, Wilk's lambda: $^{§}p = 0.018$) (n = 7).

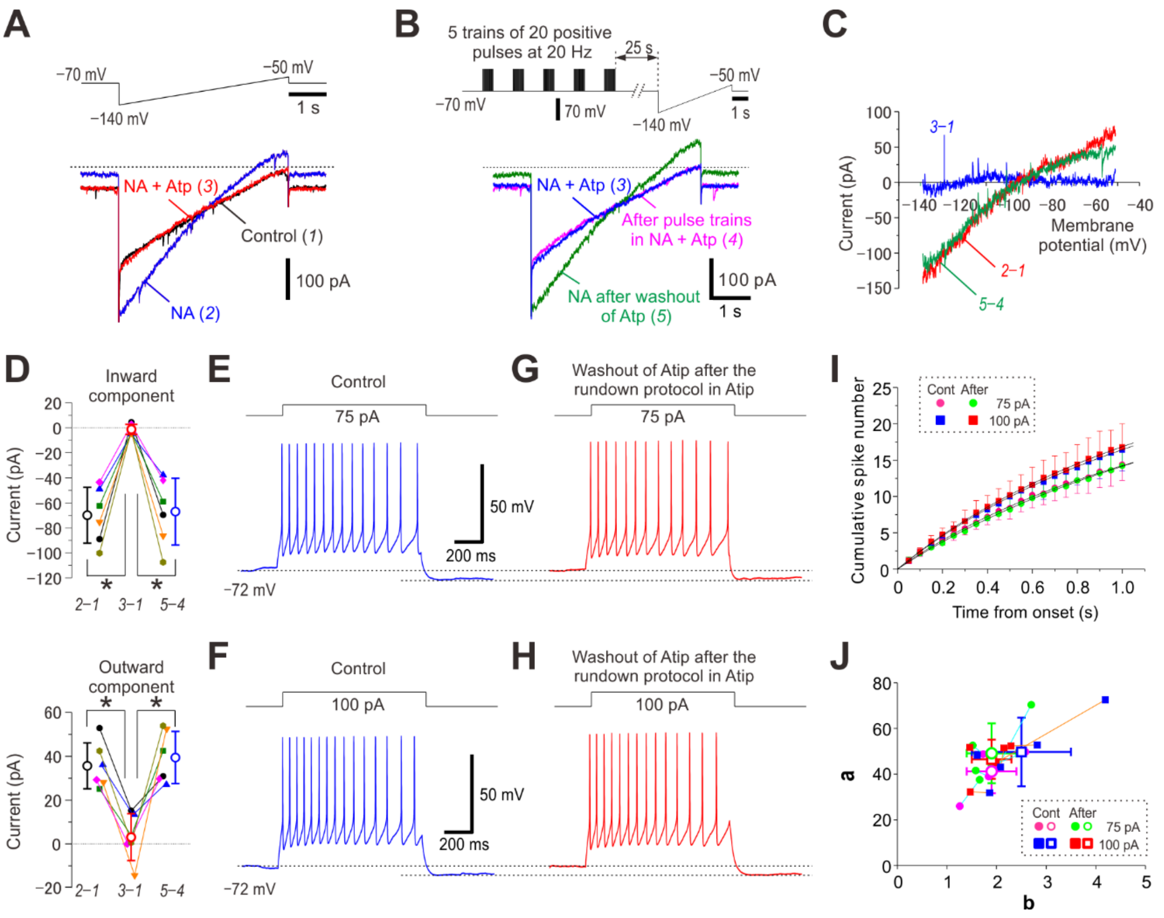


**Figure S5. Atipamezole blocks rundown of NA-induced GIRK currents, waning of spike-frequency adaptation and suppression of pulse-ADP caused by repetitive application of positive pulse trains.**

(**A**) Upper panel, Voltage command pulse. Lower panel, Superimposed current traces obtained under control condition (1), after application of 100 µM NA for 5 min (2) and after application of NA and 10 µM atipamezole for 5 min (3).

(**B**) Upper panel, A combined command pulse applied every minute; five trains of 20 positive pulses (5 ms duration to 0 mV at 20 Hz) at an inter-train interval of 2 s in the presence of extracellular 30 mM TEA, NA, atipamezole and intracellular 0.2 mM EGTA, which were followed by the ramp pulse after an interval of 25 s. Lower panel, Superimposed current traces obtained before and after 20 times application of positive pulse trains in the presence of NA and atipamezole (3 and 4, respectively) and after washout of atipamezole but still in the presence of NA (5).

(**C**) The I-V relationship of NA-induced GIRK currents obtained by subtraction of the currents recorded under the control condition from those recorded after application of NA for 5 min (red trace, 2–1), that obtained by subtraction of the control current from those recorded after application of atipamezole in addition to NA (blue trace, 3–1), and that obtained by subtraction of the currents recorded after 20 times application of positive pulse trains in the presence of NA and atipamezole from those recorded 10 min after washout of atipamezole but still in the presence of NA (green trace, 5–4).

(**D**) Pooled data showing protective effects of atipamezole on the rundown of GIRK currents; amplitudes of inward components at –130 mV and those of outward components at –60 mV at respective conditions (2–1, 3–1 and 5–4). Inward component, one-way RM ANOVA, $*p < 0.001$; outward component, $*p < 0.001$ ($n = 6$).
(**E-H**) Sample traces of spike trains induced in an LC neuron evoked by current pulses at 75 and 100 pA under control conditions and after repetitive application of positive pulse trains in the presence of atipamezole under voltage-clamp condition as shown in **A-C**. Note that atipamezole prevented the waning of spike-frequency adaptation and reduction of pulse-AHP which could have been caused by $[Ca^{2+}]_i$ increases in response to repetitive application of positive pulse trains.
(**I**) A plot of the cumulative spike numbers vs the elapsed time during the current pulses every 50 ms obtained under control condition (pink circles: 75 pA; blue squares: 100 pA) and after repetitive application of positive pulse trains in the presence of atipamezole (green circles: 75 pA; red squares: 100 pA). 75 pA current pulse, Two-way RM ANOVA, $p = 0.181$; 100 pA current pulse, Two-way RM ANOVA, $p = 0.958$ ($n = 5$).
(**J**) A plot of the saturation level (***a***) vs the half saturation constant (***b***), which were measured by curve fitting to the data points in **i**. The values of ***a*** and ***b***, which were obtained after repetitive application of positive pulse trains in the presence of atipamezole (pink circles, 75 pA; blue squares, 100 pA) were not significantly different from those obtained under control condition (green circles, 75 pA; red squares, 100 pA) (75 pA current pulse, paired *t*-test, ***a*** and ***b***, $p = 0.963$ and $p = 0.236$, respectively; 100 pA current pulse, paired *t*-test, ***a*** and ***b***, $p = 0.188$ and $p = 0.645$, respectively) and there was no significant difference in the relationship between ***a*** and ***b*** (75 pA current pulse, Wilk's lambda, $p = 0.276$; 100 pA current pulse, Wilk's lambda, $p = 0.340$) ($n = 5$).

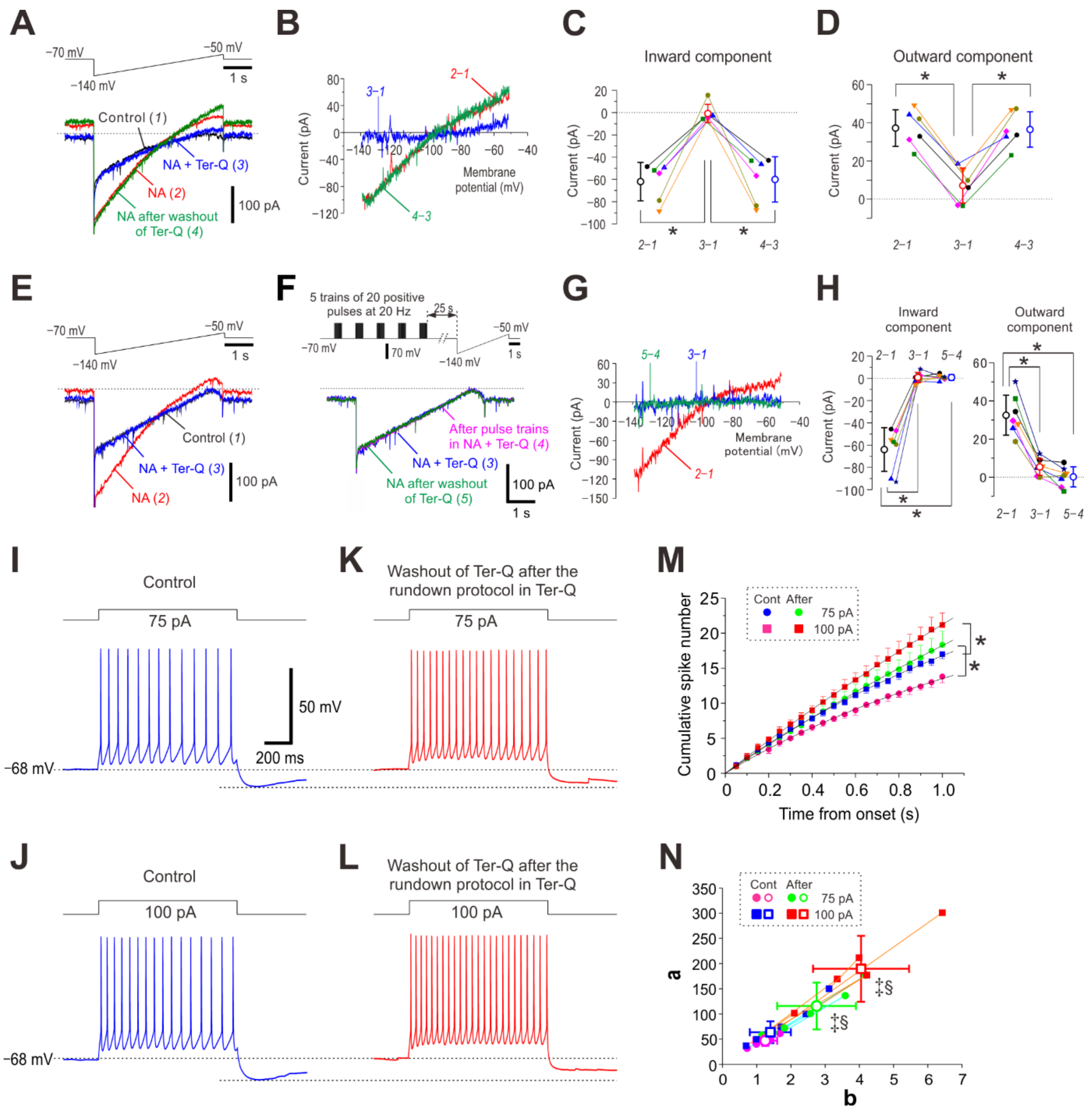

**Figure S6. Tertiapin-Q did not block $Ca^{2+}$-dependent rundown of NA-induced GIRK currents**

(**A**) Upper panel, Voltage command pulse. Lower panel, Superimposed current traces obtained under control condition (1), after application of 100 μM NA for 5 min (2), after application of NA and 200 nM tertiapin-Q (Ter-Q) (3), and after application of NA following washout of Ter-Q (4). (**B**) The I-V relationship of NA-induced GIRK-I obtained by subtraction of currents recorded under control condition from those recorded after application of NA for 5 min (red trace, 2–1), that obtained by subtraction of the control current from those recorded after application of Ter-Q in the presence of NA (blue trace, 3–1), and that obtained in the presence of NA by subtraction of currents recorded after application of Ter-Q from those recorded after washout of Ter-Q (green trace, 4–3). (**C, D**) Pooled data showing that Ter-Q reversibly decreased the amplitudes of inward components at –130 mV and those of outward components at –60 mV at respective conditions (2–1, 3–1 and 4–3) (n = 6). Inward component, one-way

RM ANOVA, *$p < 0.001$; outward component, one-way RM ANOVA, *$p < 0.001$. (**E**) Upper panel, Voltage command pulse. Lower panel, Superimposed current traces obtained under control condition (1), after application of NA for 5 min (2) and after application of NA and Ter-Q for 5 min (3). (**F**) Upper panel, A combined command pulse applied every minute; five trains of 20 positive pulses (5 ms duration to 0 mV at 20 Hz) at an inter-train interval of 2 s in the presence of extracellular 30 mM TEA, NA, Ter-Q and intracellular 0.2 mM EGTA, which were followed by the ramp pulse after an interval of 19 s. Lower panel, Superimposed current traces obtained before and after 20 times application of positive pulse trains in the presence of NA and Ter-Q (3 and 4, respectively) and after washout of Ter-Q but still in the presence of NA (5). (**G**) The I-V relationship of NA-induced GIRK-I obtained by subtraction of currents recorded under the control condition from those recorded after application of NA for 5 min (red trace, 2–1), that obtained by subtraction of the control current from those recorded after application of NA and Ter-Q (blue trace, 3–1), and that obtained by subtraction of the currents recorded after 20 times application of positive pulse trains in the presence of NA and Ter-Q from those recorded 10 min after washout of Ter-Q but still in the presence of NA (green trace, 5–4). (**H**) Pooled data showing no protective effects of Ter-Q on the rundown of GIRK-I; amplitudes of inward components at –130 mV and those of outward components at –60 mV at respective conditions (2–1, 3–1 and 5–4) (n = 7). Inward component, one-way RM ANOVA, *$p < 0.001$; outward component, one-way RM ANOVA, *$p < 0.001$. (**I-L**) Sample traces of spike trains induced in an LC neuron evoked by current pulses at 75 and 100 pA under control conditions and after washout of Ter-Q after the rundown protocol in Ter-Q as shown in **E-H**. Note the abolishment of spike-frequency adaptation and reduction of pulse-AHP after rundown of GIRK currents. (**M**) Plotting of the cumulative spike numbers vs the elapsed time during current pulses every 50 ms obtained under control condition (blue circles, 75 pA; pink squares, 100 pA) and those obtained after rundown of GIRK currents (green circles, 75 pA; red squares, 100 pA) (n = 6). 75 pA current pulse, two-way RM ANOVA, *$p < 0.001$; 100 pA current pulse, two-way RM ANOVA, *$p < 0.001$. (**N**) Plotting of the saturation level (***a***) vs the half saturation constant (***b***), which were measured by curve fitting to the data points in **M**. The values of ***a*** and ***b***, which were measured by curve fitting to the data points obtained after rundown of GIRK currents (green circles, 75 pA; red squares, 100 pA) were significantly larger than those obtained under control condition (pink circles, 75 pA; blue squares, 100 pA) (75 pA current pulse, paired *t*-test, ***a*** and ***b***, $^{‡}p = 0.015$ and $^{‡}p = 0.010$, respectively; 100 pA current pulse, paired *t*-test, ***a*** and ***b***, $^{‡}p = 0.004$ and $^{‡}p < 0.001$, respectively, and there was a significant difference in the relationship between ***a*** and ***b*** (75 pA current pulse, Wilk's lambda: $^{§}p = 0.020$; 100 pA current pulse, Wilk's lambda: $^{§}p = 0.010$) (n = 6).

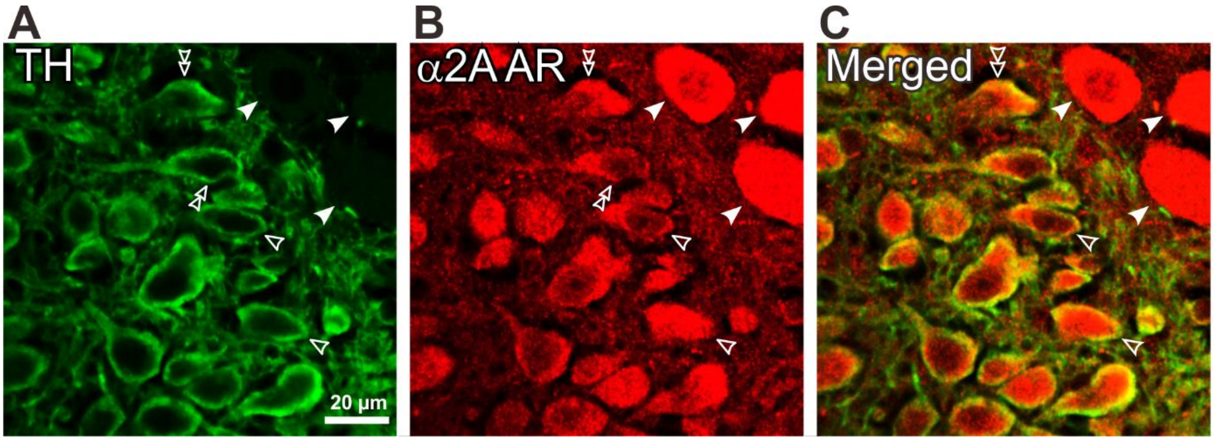


**Figure S7. Reliability of anti-α2A-AR antibody revealed by differential expression of α2A ARs between LC and MTN.**

(**A-C**) Confocal images of LC and MTN neurons showing the immunoreactivities for TH and α2A ARs, together with a merged one. Filled arrowheads indicate MTN neurons, in which TH was not expressed, but α2A ARs were more extensively expressed than in LC neurons. Open arrowheads indicate oval shaped neurons. Open double arrowheads indicate rhombus-like shaped neurons. Differential staining of α2A-ARs between MTN and LC neurons verifies the reliability of the antibody against α2A-ARs.

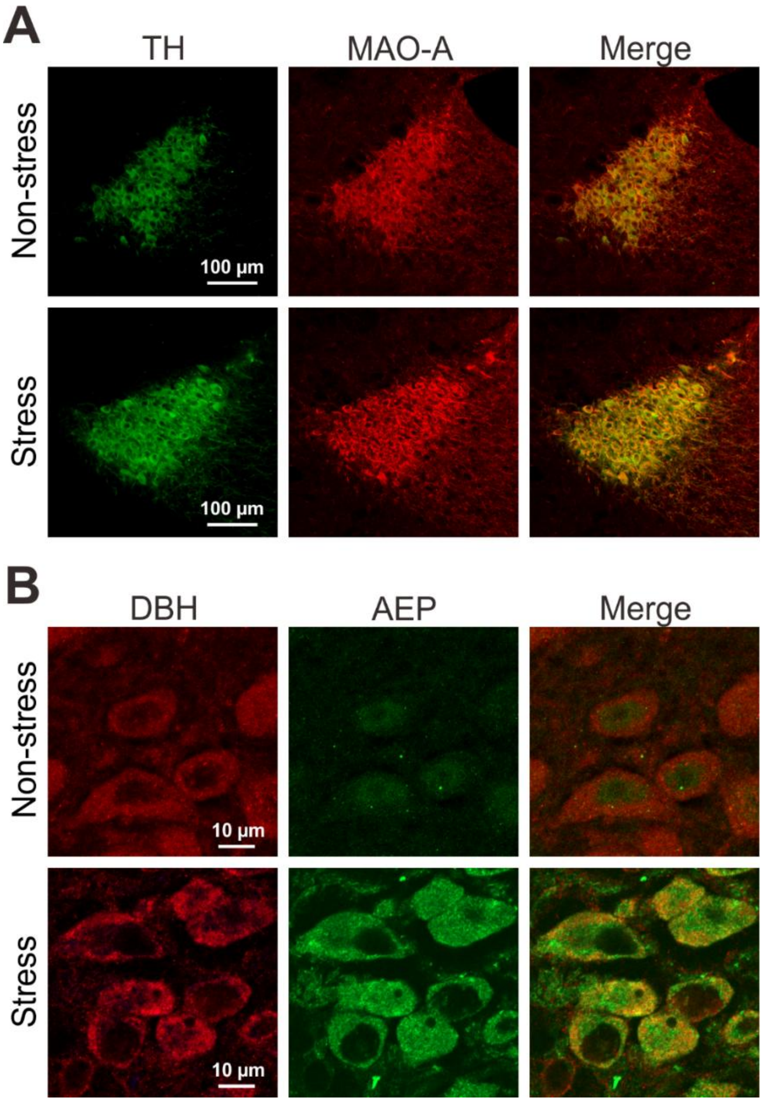


**Figure S8. Immunoreactivities of TH, MAO-A, DBH and AEP in LC neurons.**

(**A**) Confocal images of neurons in LC at a low magnification showing the immunoreactivities for TH and MAO-A, together with a merged one. RS appeared to cause slight increases in the expressions of TH and MAO-A. Brightness of background in sample image obtained from the control was corrected to be equal to that obtained from the RS.

(**B**) Confocal images of LC neurons at a high magnification showing the immunoreactivities for DBH and AEP, together with a merged one. RS caused increases in the expression of AEP.

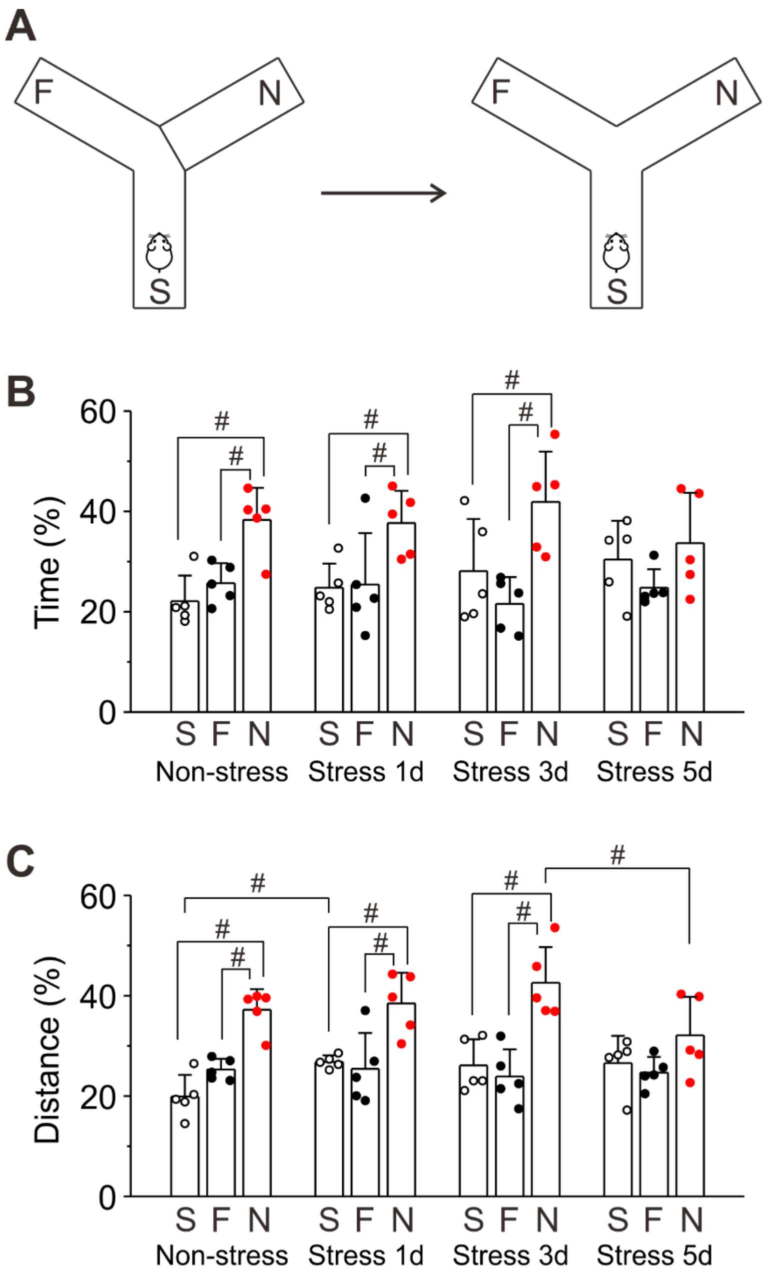


**Figure S9. RS-induced impairment of spatial memory.**

(**A**) Left panel: Y maze to explore 2 arms, the start (S) and familial (F) arms with the novel (N) arm closed, for 15 minutes of training. Right panel: Y maze to explore 3 arms (the S, F and N arms) for a 5-minute test session.

(**B**) Time spent in S, F, and N arms in non-stress mice and in 1-day, 3-day and 5-day RS mice. Two-way RM ANOVA, #$p < 0.05$.

(**C**) Distance traveled in S, F, and N arms in non-stress mice and in 1-day, 3-day and 5-day RS mice. Two-way RM ANOVA, #$p < 0.05$.

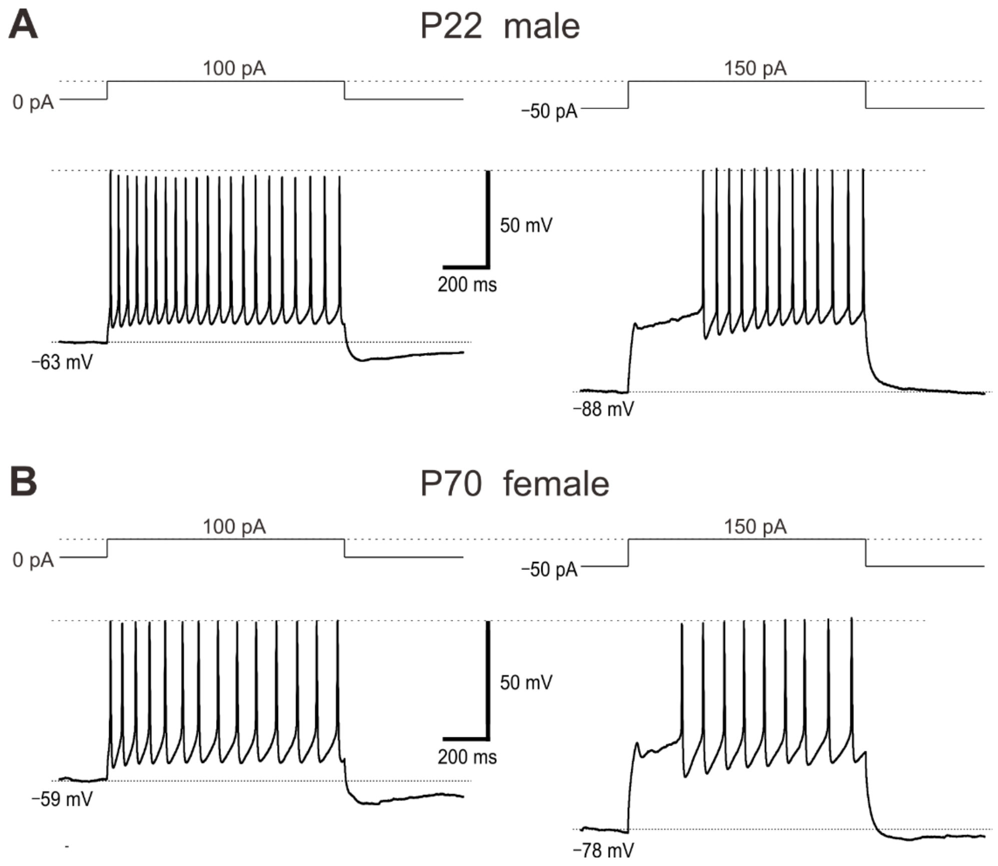


**Figure S10. Spike-frequency adaptation and A-like $K^+$ current in a male juvenile mouse and a female adult mouse.**

(**A**) An LC neuron of a male mouse at postnatal days 22 (P22) displayed spike-frequency adaption in response to a 100 pA current pulse at –63 mV while it displayed a late spiking due to the presence of A-like $K^+$ current in response to a 150 pA current pulse at –88 mV. (**B**) An LC neuron of a female mouse at P70 displayed spike-frequency adaption in response to a 100 pA current pulse at –59 mV while it displayed a late spiking due to the presence of A-like $K^+$ current in response to a 150 pA current pulse at –78 mV. There were no apparent differences in spike-frequency adaptation and A-like K+ current between juvenile and adult mice and also between male and female mice.
